# Comparative analysis of uncoupled succinate production by the Fe^II^/2-oxoglutarate-dependent dioxygenases

**DOI:** 10.1101/2024.09.09.612061

**Authors:** Susmita Das, Saumya Ranjan, Carmel Keerthana, Gayathri Seenivasan, Nikhil Tuti, Unnikrishnan P Shaji, Gargi Meur, Roy Anindya

**Author notes:** Corresponding author: Roy Anindya.

## Abstract

Non-heme iron (Fe^II^) and 2-oxoglutarate(2OG)-dependent dioxygenases catalyse a diverse array of biological reactions. These enzymes couple the oxidative decarboxylation of 2OG to the hydroxylation of the substrates. However, in the absence of the substrate, oxidative decarboxylation of 2OG generates succinate. We have determined succinate level by using succinyl-CoA synthetase to monitor this uncoupled decarboxylation of Fe^II^/2OG-dependent dioxygenases and measured the uncoupled 2OG turnover of different Fe^II^/2OG-dependent dioxygenases. We also performed comparative analysis and verified the functionality of human dioxygenase ALKBH6.

## 1. Introduction

Iron and 2-oxoglutarate (2OG)-dependent dioxygenases (Fe^II^/2OG dioxygenase) are involved in a variety of biochemical reactions, such as chromatin modification, fatty acid metabolism, regulating the hypoxic response and repair of alkylated DNA [1]. These enzymes utilize non-haem ferrous iron (Fe^II^) as a co-factor and 2OG as a co-substrate [2]. Oxidative decarboxylation of 2OG leads to the release of CO_2_ and the formation of succinate [3]. The reaction catalysed by Fe^II^/2OG-dependent dioxygenases consists of two half-reactions. The first half-reaction is O_2_-activation leading to oxidative decarboxylation of 2OG to succinate and CO_2_. The second half-reaction involves the transfer of the activated oxygen to the prime substrate, resulting in diverse outcomes such as desaturation, demethylations, ring-closure, and hydroxylation. These two half-reactions are generally coupled, however, sometimes decarboxylation of 2OG takes place even in the absence of any substrate. Such 2OG turnover is termed an uncoupled reaction and generally results in the oxidation of catalytic ferrous iron (Fe^II^) which is subsequently reversed by ascorbic acid. The uncoupled reaction was described in many Fe^II^/2OG-dependent dioxygenases, including prolyl hydroxylase, lysyl hydroxylase and asparaginyl hydroxylase [4-6] and the AlkB family of nucleic acid demethylase [7], including ALKBH2, ALKBH3 [8], ALKBH5 [9] and ALKBH8 [10]. The rate of this uncoupled reaction was observed to be approximately 1-5% of the rate in the presence of substrate [11]. Due to distinct substrate specificity, it is not straightforward to compare the efficiency of Fe^II^/2OG dioxygenases and we sought to examine if the comparative analysis of uncoupled 2OG turnover can be performed by determining the uncoupled activity. The Fe^II^/2OG dioxygenase superfamily is extremely diverse and includes many enzymes without any known substrate. We were also interested to know whether the uncoupled conversion of 2OG to succinate can be used as a surrogate reaction to validate their functionality.

Most of the reports on the uncoupled activity of Fe^II^/2OG-dependent dioxygenase involved the detection of the product released by adding ^14^C-labeled 2OG and resulting 14C-labelled succinate formation measured by scintillation counting [12]. Mass spectrometry-based methods for directly monitoring the product or the substrate were also applied [13, 14]. Because of the difficulty of mass spectrometry or radiometric assay, we used a colorimetric method for measuring uncoupled activity that employs *E coli* succinyl-CoA synthetase enzyme (SucCD) [15, 16]. In this report, we demonstrate the functionality of an AlkB family of Fe^II^/2OG-dependent dioxygenase, the specific substrate of which remains unidentified. We also illustrate that succinate detection by this method is suitable for comparing the uncoupled activity of some well-characterized human Fe^II^/2OG-dependent dioxygenases involved in DNA repair.

## 2. Materials and Methods

### 2.1 Standard curve

A standard curve of phosphate was prepared using increasing concentrations of NaH_2_PO_4_ (0-200 μM) in 20 mM MES buffer, pH 6.5 and mixed with 100 μL of colorimetric solution (1:1 vol) consisting of vanadate-molybdate reagent. The mixture (200 μL) was incubated at room temperature for 5 min. Succinate standard curve was prepared by using increasing concentrations (0-200 μM) of succinate in buffer (20 mM MES buffer, pH 6.5). The colorimetric solution (100 μL) was added (1:1 vol) and the mixture (200 μL) was incubated at room temperature for 5 min. All the absorbance was recorded at 660 nm using a 96-well plate reader (Molecular Devices SpectraMax M5).

### 2.2 Cloning, expression and purification of recombinant *E coli* SucCD protein

*E coli* succinyl-CoA synthetase is a heterotetramer composed of *sucC* and *sucD* genes. The *sucC* gene encodes a polypeptide of 41.4 kDa that corresponds to the IZ subunit of succinyl-CoA synthetase. The *sucD* gene encodes a 29.7 kDa polypeptide, which constitutes the CI subunit of succinyl-CoA synthetase. The sucC and sucD genes are translationally coupled as the stop codon present following the sucC gene overlaps with the sucD initiation codon by a single nucleotide [17]. Therefore, *sucC* and *sucD* genes were amplified by PCR using specific forward and reverse primers corresponding to sucC and sucD, respectively. The PCR amplified DNA was cloned between *NdeI* and *XhoI* of pET28a such that sucC and sucD proteins have N terminal and C terminal his-tag, respectively. The SDS-PAGE analysis of His-sucCD revealed two bands, one representing 43 kDa β-subunit (His-sucC) and the other band being 31 kDa α-subunit (His-sucD). The heteromeric protein was purified by using Ni-NTA agarose followed by gel-filtration chromatography.

### 2.3 Purification of recombinant Fe^II^/2OG-dependent dioxygenase

Recombinant Fe^II^/2OG-dependent dioxygenase, including AlkB, ALKBH2, ALKBH3, and Tpa1 were cloned as described before [18, 19]. Recombinant His-tag AlkBH6 was a generous gift from Prof Timothy O’Connor, City of Hope, Duarte Cancer Centre, USA. AlkB mutant (H131A, H133A), ALKBH2 mutant (H171A, D173A), ALKBH3 mutant (H191A, D193A, H257A), ALKBH6 mutant (H114A, D116A, H182A), Tpa1 mutant (H159C, D161N, H227C, H237C, R238A) were generated by site-directed mutagenesis. Wild type and mutant recombinant proteins were purified using standard Ni-affinity followed by gel filtration chromatography. Site-directed mutagenesis was used to generate the catalytic mutants of each enzyme as reported earlier [18, 19].

### 2.4 Succinate detection

The assay was performed in 96-well plate with 200 μl reaction volume. The succinate production by Fe^II^/2OG dioxygenase (5-20 μM) were carried out in the buffer containing 20 mM Tris-HCl, pH 8.0, 2 mM ascorbate, 20 μM ammonium ferrous sulfate and 0.2 mM 2OG in final volume of 50μl. The uncoupled reaction mixture was incubated up to 1 h at 37 □C. Then the succinate formed in the reaction was detected by adding another 50 μl reaction mixture containing and Suc-CD enzyme (1 μM) in reaction buffer (20 mM MES, pH 6.5, 1 mM CoA,1 mM ATP, 10 mM MgCl_2_). The Suc-CD enzyme reaction was incubated for 1 h at 37 □C. After incubation, the final succinate values were obtained from the succinate standard curve. The assay was also performed in the presence of inhibitor (nickel chloride, 5 mM). The results were plotted and analyzed using GraphPad Prism.

### 2.5 Time-dependent succinate detection

The time-dependent succinate production was measured by incubating dioxygenase enzyme (10 μM) in reaction buffer (50 μl) at 37 □C. Part of the reaction mixture was removed every 20 min and stopped by adding 2 mM Nickel Chloride. Then sucCD was added and the reaction continued as mentioned above. The rate of the reaction was calculated from the slope of linear regression in GraphPad Prism.

## 3. Results and discussion

### 3.1 Comparative analysis of uncoupled succinate production of Fe^II^/2OG-dependent dioxygenases

In the absence of substrate, active Fe^II^/2OG-dependent dioxygenases decarboxylate the co-substrate 2OG to produce succinate and CO_2_. Succinyl-CoA synthetase (SucCD) was used to determine succinate and with the concomitant hydrolysis of ATP to form ADP and phosphate. Using a molybdate reagent, the phosphate released was quantified colorimetrically [15]. We applied the same method for comparing the uncoupled succinate formation by four different Fe^II^/2OG-dependent dioxygenases (**Figure 1A**). All the recombinant His-tag proteins were purified (**Figure 1B**) as described before [18-22]. One of the important members of the Fe^II^/2OG-dependent dioxygenase family is *E coli* DNA repair enzyme AlkB [23], which catalyses the demethylation of 1meA and 3meC from DNA and RNA substrates. When the reaction was carried out in the presence of recombinant AlkB, 2OG and ascorbate, but without any alkylated DNA substrate, the level of succinate production was determined (**Figure 1C**). A highly conserved HX(D/E)X_n_H triad motif supports acid residues responsible for Fe^II^-binding and catalysis in all Fe^II^/2OG dioxygenases [24]. To confirm that only catalytically active Fe^II^/2OG dioxygenases can perform uncoupled 2OG turnover, we evaluated catalytically inactive mutant AlkB in the absence of substrate. As shown in **Figure 1D**, this catalytically inactive AlkB did not show any uncoupled succinate production and suggested that this method could be reliably used to confirm uncoupled activity. Next, we evaluated *Saccharomyces cerevisiae* Fe^II^/2OG dioxygenase Tpa1 and its catalytic mutant [19]. We confirmed the uncoupled succinate production from wild-type Tpa1 (**Figure 1D**). We further examined two human AlkB homologues, ALKBH2 and ALKBH3. As a negative control, the succinate production of the catalytically dead ALKBH2 and ALKBH3 mutant were evaluated. As shown in **Figures 1E** and **F**, ALKBH2 and ALKBH3 displayed uncoupled activity but not the catalytic mutants. We observed that some succinate is still produced in ALKBH mutants with all the catalytic triad residues removed. This could be due to hydrolysis of ATP by some undetected contaminant present in the mutant protein fractions.

**Figure 1.**
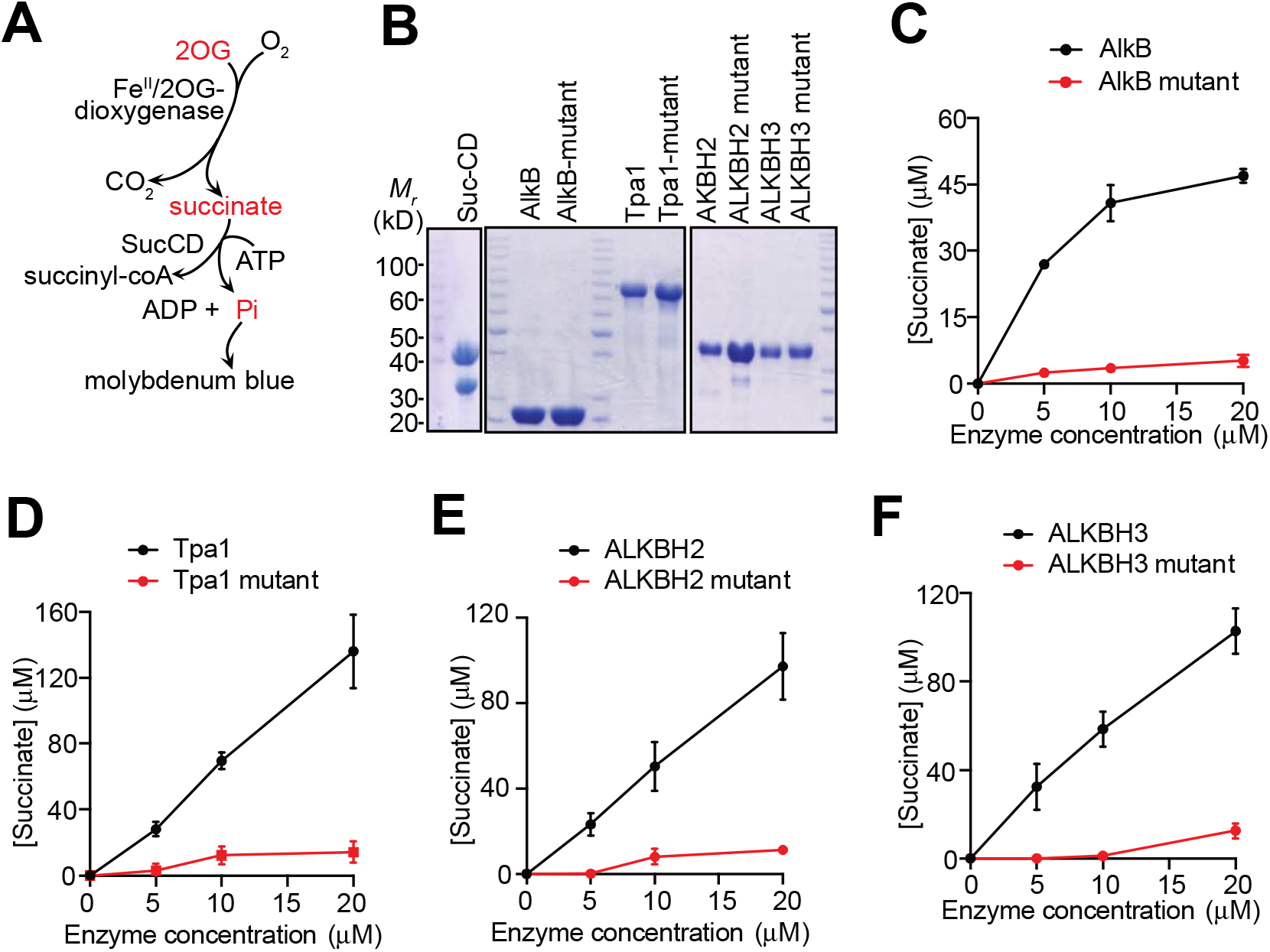
Comparative analysis of uncoupled succinate production by Fe^II^/2OG-dependent dioxygenases. **(A)** Outline of generic reaction catalysed by Fe^II^/2OG-dependent dioxygenases and SucCD enzymes **(B)** SDS-PAGE analysis of *E coli* SucCD, wildtype and mutant AlkB, Tpa1, ALKBH2, and ALKBH3. It should be noted that the protein loading is not equal for all proteins and differentially stained SDS-PAGE gels is depicted. **(C)** Enzyme concentration-dependent production of succinate monitored by the reaction of *E coli* AlkB,**(D)** *S cerevisiae* TPA1, **(E)** ALKBH2, **(F)** ALKBH3. Error bar represent mean ± S.E. (n = 4).

Among the proteins examined, AlkB showed a saturation type succinate production with increasing concentration, unlike others. Presently, we do not know if this could be due to enzyme self-inactivation or some other mechanism. Taken together, these results suggest that succinate production could be used for determining uncoupled activity of Fe^II^ or 2OG binding in structure-function studies of this family of enzymes.

### 3.2 ALKBH6 is catalytically active Fe^II^/2OG-dependent dioxygenase

Although many Fe^II^/2OG-dependent dioxygenases are encoded in the human genome, it is not known whether they are all functional proteins. One such enzyme is the human AlkB homologue ALKBH6. This enzyme was structurally and functionally characterized but the substrate of this enzyme is not known yet and biochemical activity could not be determined [26, 27]. We decided to verify the functionality of ALKBH6 using recombinant ALKBH6 and its catalytic mutant (**Figure 2A**). When an increasing concentration of ALKBH6 was incubated with the co-substrate 2OG without any substrate, succinate was produced in a dose-dependent manner (**Figure 2B**). Previous studies reported that nickel could replace the iron (Fe^II^) in the catalytic centre of the Fe^II^/2OG-dependent dioxygenases and cause inhibition of their enzymatic activity [25]. We monitored nickel-mediated inhibition of uncoupled 2OG turnover by adding nickel chloride to Fe^II^-bound proteins. As expected, it reduced succinate production for all the enzymes tested (**Figure 2C**). Interestingly, the magnitude of the inhibition varied between the enzymes, which might be due to the different affinities for the catalytic Fe^II^ among the enzymes. Next, we used nickel-mediated inhibition to block 2OG production at different time points to measure time-dependent succinate production and derived the rate of uncoupled reaction (**Figure 2D**). We observed that ALKBH3 showed highest rate of uncoupled succinate turnover (1.02 µM min^-1^) compared to ALKBH2 (0.46 µM min^-1^) and ALKBH6 (0.37 µM min^-1^). These results indicate that ALKBH6 is functionally active Fe^II^/2OG-dependent dioxygenase and might display robust activity when the right substrate is identified.

**Figure 2.**
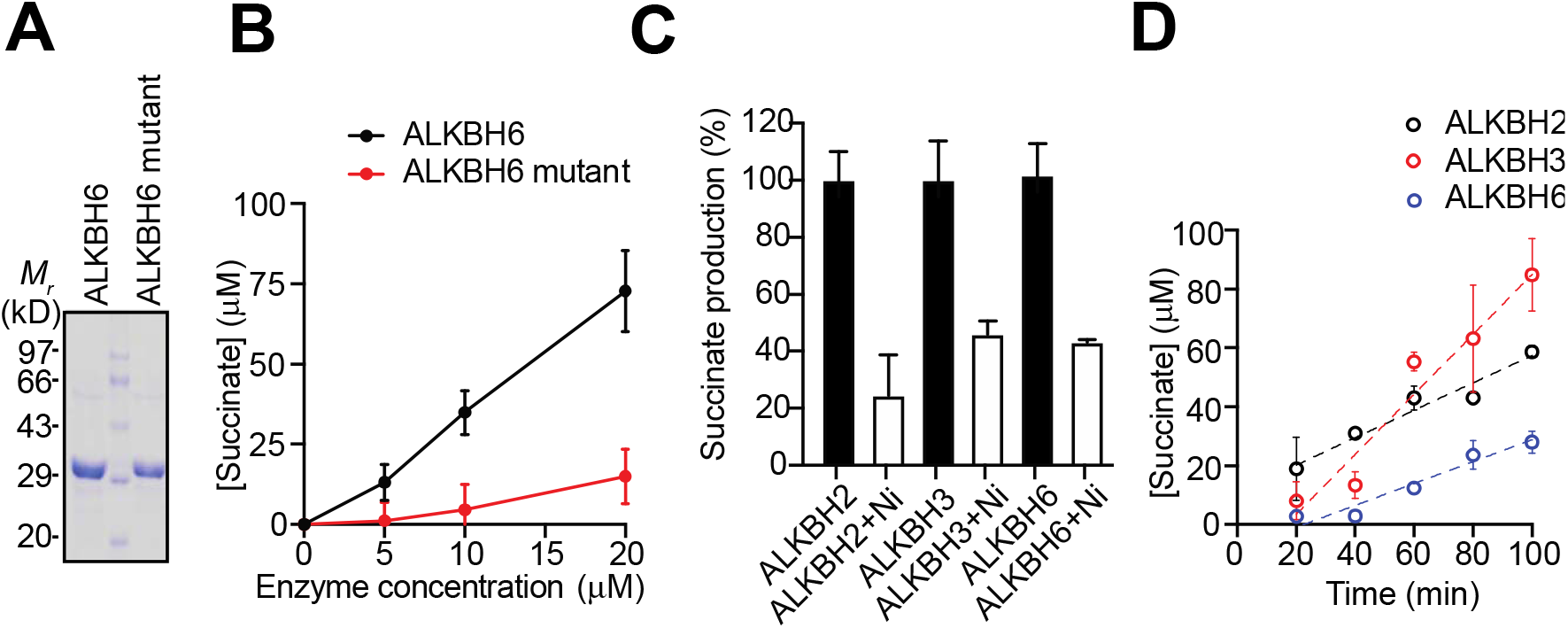
Production of uncoupled succinate by ALKBH6. **(A)** SDS-PAGE analysis of wildtype and mutant ALKBH6. **(B)** Enzyme concentration-dependent production of succinate. (**C**) Inhibition of succinate production in the presence of nickel (Ni) ions. (**D**) Comparison of time-dependent increase of succinate production by ALKBH2, ALKBH3 and ALKBH6 (10 μM). Dashed lines represent the linear regression used for calculating the rate of succinate production. Error bar represent mean ± S.E. (n = 4).

## Conclusion

We demonstrated that the uncoupled succinate production could be used for comparing the uncoupled activity of the Fe^II^/2OG-dependent dioxygenases and establishing the functionality of any Fe^II^/2OG-dependent dioxygenases when the identity of the substrate is unknown.

